# Addressing the reliability fallacy: Similar group effects may arise from unreliable individual effects

**DOI:** 10.1101/215053

**Authors:** Juliane H. Fröhner, Vanessa Teckentrup, Michael N. Smolka, Nils B. Kroemer

**Author notes:** **Corresponding authors:** Juliane Fröhner, M.Sc., Section of Systems Neuroscience, Department of Psychiatry and Psychotherapy, Technische Universität Dresden, Würzburger Str. 35, 01187 Dresden, Germany, E-Mail, Nils B. Kroemer, Dr., Department of General Psychiatry and Psychotherapy, University of Tübingen, Calwerstr. 14, 72076 Tübingen, Germany.

## Abstract

To cast valid predictions of future behavior or diagnose disorders, the reliable measurement of a “biomarker” such as the brain activation to prospective reward is a prerequisite. Surprisingly, only a small fraction of functional magnetic resonance imaging (fMRI) studies report or cite the reliability of brain activation maps involved in group analyses. Here, using simulations and exemplary longitudinal data of 126 healthy adolescents performing an intertemporal choice task, we demonstrate that reproducing a group activation map over time is not a sufficient indication of reliable measurements at the individual level. Instead, selecting regions based on significant main effects at the group level may yield estimates that fail to reliably capture individual variance in the subjective evaluation of an offer. Collectively, our results call for more attention on the reliability of supposed biomarkers at the level of the individual. Thus, caution is warranted in employing brain activation patterns prematurely for clinical applications such as diagnosis or tailored interventions before their reliability has been conclusively established by large-scale studies. To facilitate assessing and reporting of the reliability of fMRI contrasts in future studies, we provide a toolbox that incorporates common measures of global and local reliability.

## Introduction

Since the early 1990s, researchers use functional magnetic resonance imaging (fMRI) to characterize general aspects of brain function which are immutable (or “fixed”) within a population. Hence, many paradigms were optimized for low between-subject variability (Hedge, Powell, & Sumner, 2017) typically leading to strong main effects in analyses at the group level. However, the advent of the Research Domain Criteria (RDoC) has led to a surge of interest in individual “biomarkers” for mental disorders (Insel et al., 2010). Nevertheless, the investigation of *intra-individual* variability and stability is still a relatively young, but quickly growing field in fMRI research (Dubois & Adolphs, 2016; Garrett et al., 2013; Kroemer et al., 2016; Van Horn, Grafton, & Miller, 2008; Vetter et al., 2017). One of the key challenges is to identify an appropriate mapping between individual brain activation and behavior (Finn et al., 2017). A prerequisite for this endeavor is to formally establish that a proposed biomarker, supposed to capture individual neurobiological characteristics of brain function (Insel et al., 2010), can indeed be reliably measured to predict future behavior.

To define such basic statistical requirements for a candidate biomarker, key benchmarks for reliability have been previously established in individual differences research (Dubois & Adolphs, 2016). Reliability is a prerequisite for measurements to be ultimately valid, but it is not well known what the reliability of fMRI brain activation is, even for popular paradigms in the literature (Bennett & Miller, 2010; Vul, Harris, Winkielman, & Pashler, 2009). Given the variety of studies and study designs, it is pivotal to evaluate and report reliability for each scenario. Reliability is critically important when the scientific objective is to predict or classify as it is often the case in longitudinal or clinical studies (Dubois & Adolphs, 2016). Already in the beginning of the last century, Spearman (1910) pointed out that it is harder to distinguish between persons by a less reliable measure, making it harder to detect associations with other constructs as a result. This raises the question to what extent fMRI brain responses could be used to predict treatment outcomes when they are not reliably measured within patients in the first place (Nord, Gray, Charpentier, Robinson, & Roiser, 2017). Arguably, not every biomarker must be stable over time to be of clinical use, for example if it reflects an acute state of a disorder. Nevertheless, reliability is mandatory for any risk factor that confers liability and is intended to predict the onset and etiology of a disorder.

Psychological measures are commonly regarded as reliable when their reliability exceeds 0.8 (Cicchetti & Sparrow, 1981; Cicchetti, 2001), which is hardly achieved (Hedge et al., 2017). Illustratively, Hedge et al. (2017) have highlighted the antagonism between maximizing robust group-level effects on the one hand (“fixed effect”) and reliably detecting individual differences on the other hand (“random effect”). Classical experimental research aims to minimize inter-individual variability by identifying robust effects at the group level and, ideally, in every individual. In contrast, individualized prediction is critically dependent on reproducible differences between individuals, which are captured by random effects in statistical models. Consequently, there is a trade-off between optimizing within-subject or between-subject effects, because they represent independent sources of variance and count as error in the analysis of the other (Yarkoni & Braver, 2010).

With respect to fMRI, generalizable effects were of main interest for a long time. Therefore, researchers focused initially on the reliability of fMRI at the group level (Aron, Gluck, & Poldrack, 2006; Fliessbach et al., 2010; Gee et al., 2015). Still, Paul et al. demonstrated recently that even large fMRI studies (i.e. N = 100) do not produce group results with good reliability (Paul, Turner, Miller, & Barbey, 2017). Furthermore, previous studies showed that group-level stability is not indicative of individual stability (Raemaekers et al., 2007; van den Bulk et al., 2013; Vetter, Pilhatsch, Weigelt, Ripke, & Smolka, 2015), while research focusing on within-subject reliability over time has produced mixed results. Good reliability was shown during performance monitoring in adolescents and adults (Koolschijn, Schel, Rooij, Rombouts, & Crone, 2011). Moreover, Plichta et al. (2012) reported differential within-subject reliability for three tasks with similarly high between-subject reliability. For two tasks (motivational and cognitive), within-subject reliability was fair to good, whereas reliability was low for the emotional task (Plichta et al., 2012), which is in line with the low reliability in an emotional face processing task (Nord et al., in press). Recently, our group analyzed reliability for three different tasks including a subset of the data reported here (Vetter et al., 2017). They showed good reliability in an emotional attention and an intertemporal choice task, but only fair reliability for a cognitive control task (Vetter et al., 2017). However, their analysis focused on conditions contrasted against baseline and not on parametric or difference contrasts, which are commonly used (Bickel, Pitcock, Yi, & Angtuaco, 2009; Hare, Camerer, & Rangel, 2009; Kable & Glimcher, 2007; Wittmann, Lovero, Lane, & Paulus, 2010) and thus need to be evaluated regarding their reliability as well.

To sum it up, even though the reliability of data is a prerequisite in studying individual differences and fMRI brain activation is increasingly applied to his end, little is known about fMRI reliability to date. A straightforward answer to the simple question on the reliability of a given paradigm is further complicated by considerable variability in the existing literature in terms of analysis level (group vs. individual) and reliability measures employed (local measures: e.g., intraclass correlation coefficients (ICC) vs. global measures: e.g., overlap coefficients). Moreover, cross-sectional reliability has received surprisingly little attention so far since low longitudinal reliability might arise from different sources of error (e.g., state effects). Here, we are providing analyses of cross-sectional and longitudinal reliability in simulated data and in a sample of adolescents investigated during an intertemporal choice task at the age of 14, 16 and 18. To facilitate future comprehensive assessment of fMRI reliability, we introduce the collection of the employed global and local measures, which will be bundled in the MATLAB toolbox *fMRelI*.

## Materials and methods

### Longitudinal data case: delay discounting

To illustrate the assessment of reliability of fMRI contrast maps, we will use a previously reported longitudinal study. Briefly, adolescents were repeatedly investigated at the ages of 14, 16 and 18, mainly to examine the influence of substance consumption on brain development (for details see Jurk, Mennigen, Goschke, & Smolka, 2016; Ripke et al., 2012, 2014; Rodehacke et al., 2014; Vetter et al., 2015). Recently, we investigated the reliability of selected fMRI contrasts for the first two acquisition waves (Vetter et al., 2017). Vetter et al. (2017) looked at three different tasks using the between-session intraclass correlation coefficient focusing on simple contrasts against baseline. Here, the aim is to provide a substantially extended analysis for various within- and between-session measures of group-level and individual stability using an exemplary paradigm, which is why we focus on the intertemporal choice paradigm across all three acquisition waves.

#### 1. Participants

The presented data originate from the project “the adolescent brain”. To prospectively study brain function and substance use, participants were recruited at the age of 14 years and re-invited at the ages of 16 and 18. During the three acquisition waves, participants underwent an extensive assessment, including fMRI sessions and an intertemporal choice task (Ripke et al., 2012, 2014). Initially, 250 adolescents participated in the study. In total, 151 of them completed the task during every acquisition wave. For the reliability analysis, we excluded eight participants due to a diagnosis of a mental disorder because the onset might distort individual reliability of a candidate biomarkers. Additionally, 17 participants were excluded because they had more than 10% invalid trials in at least one session. Invalid trials were defined as missing or implausible responses such as deciding for a reward with a subjective value lower than half of the alternative reward (see Ripke et al., 2012). This criterion was imposed to exclude all participants who were not sufficiently attentive or internally consistent in their decisions. Thereby, 126 individuals remained for the reliability analysis.

#### 2. Functional Magnetic Resonance Imaging

##### Paradigm

During the intertemporal choice task, participants choose between a smaller immediate amount of money and a larger delayed amount. To balance choices for immediate or delayed offers, the offers during the fMRI session are adapted to the individual discounting rate, *k*, determined during a training session at age 14. The temporal discounting rate, *k*, governs the subjective assignment of value, *V*, to a monetary amount *A* when it is delivered after delay *D*:

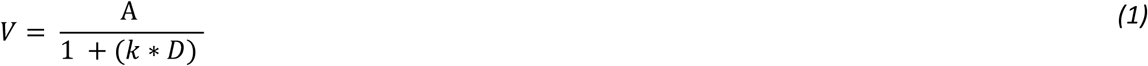

During quality control, we identified that the manually entered discount rate was not the same in at least one session for five subjects. However, the effects on the adaptation of the task were minor and results did not change after excluding these subjects, which is why they were retained in the current analysis.

##### fMRI data acquisition and analysis

Functional data was acquired with a 3 T whole-body MR tomograph (Magnetom TRIO, Siemens, Erlangen, Germany) equipped with a 12-channel head coil at the Neuroimaging Centre Dresden. A standard echo planar imaging (EPI) sequence was used for the functional images [repetition time (TR): 2410ms; echo time (TE): 25ms; flip angle: 801°; number of slices: 42; slice thickness: 2mm (1mm gap); field of view (FoV): 192×192mm^2^; resampled to voxel size: 3×3×3mm^3^]. For structural images, a 3D T1-weighted magnetisation-prepared rapid gradient echo (MPRAGE) image was acquired (TR: 1900 ms, TE: 2.26 ms, FOV: 256×256 mm^2^, 176 slices, voxel size: 1×1×1mm^3^, flip angle: 91; for details, see Ripke et al., 2012).

fMRI data analysis was performed using SPM12 (Wellcome Department of Neuroimaging, London, United Kingdom) and MATLAB R2015a (Mathworks, Inc., Sherborn, MA). The preprocessing followed a standard pipeline including slice-time correction, realignment, coregistration to the respective structural image of the participant, normalization to the standard EPI template [Montreal Neurological Institute (MNI)] and smoothing with an isotropic Gaussian kernel (8mm full-width at half-maximum). The first-level regressors included one regressor representing the offer onset and the corresponding parametric modulator. The parameter represents the subjective value of the presented offer, which was calculated via *Equation 1* using the *k* determined at age 14. We employed the same discount rate for all three waves because we found that there was no significant change overall and using a single *k* value to calculate subjective value improves comparison across waves. Please note though that minor changes in *k* only have negligible effects on the estimated subjective value and parametric regressors would be highly correlated in any case. At the end of each trial, an exclamation mark appeared at one side of the screen, indicating where participants had to press to select the presented (delayed) offer. To separate the corresponding motor responses, we included two regressors representing the onsets of button presses with the left and right hand, respectively. We included the six realignment parameters as nuisance regressors. In line with previous research, we focused primarily on the offer and the subjective-value contrasts. Nevertheless, we also report the reliability of the motor contrasts, which have been previously shown to be good in terms of retest reliability (Havel et al., 2006; Loubinoux et al., 2001; Marshall et al., 2004; Waldvogel, van Gelderen, Immisch, Pfeiffer, & Hallett, 2000).

### Simulation

First, we validated the toolbox using simulated data that originated from the first wave. Therefore, we used the contrast maps for participants at the age of 14 years and simulated longitudinal changes in contrast maps at the age of 16 and 18, respectively. We simulated different levels of intra-individual stability of the primary contrast of interest ‘subjective value’ over time (mean parameter value = 0.008 and standard deviation = 0.02) using a known range of random Gaussian noise (sigma: 0.01 - 0.04) and compared similarity matrices for the simulated and the actual data.

### Analysis workflow for reliability estimation

To illustrate the workflow in the assessment of reliability at different levels, we will describe the cross-sectional and longitudinal analyses of reliability via various measures implemented in the toolbox fMRelI (see *Figure 1*). fMRelI is an open-access toolbox, available via github (<u>https://github.com/nkroemer/reliability</u>). It is a MATLAB-based graphical user interface (GUI). The toolbox requires SPM12 and the “Tools for NIfTI and ANALYZE image” (<u>https://de.mathworks.com/matlabcentral/fileexchange/8797-tools-for-nifti-and-analyze-image</u>).

**Figure 1:**
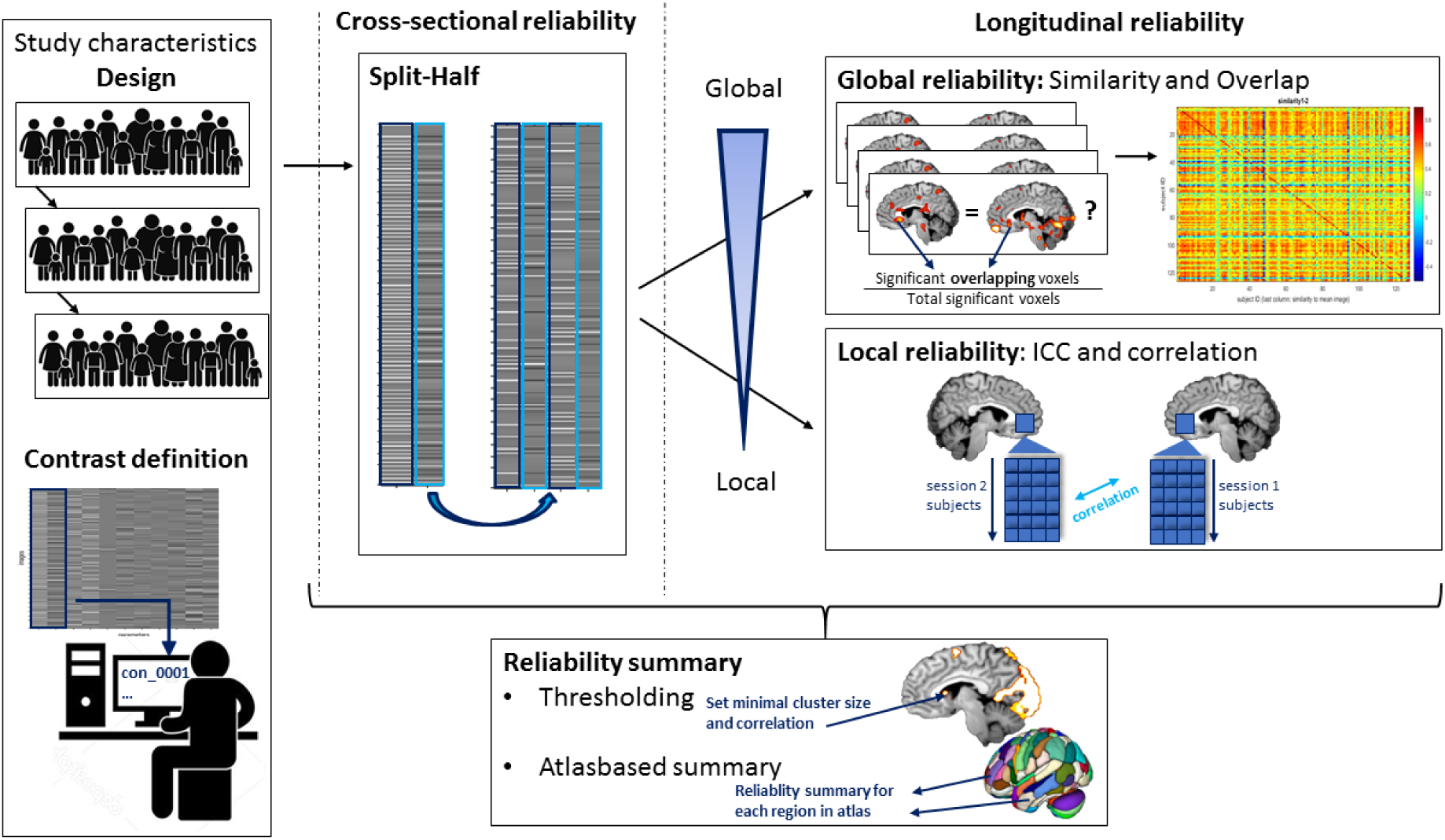
Schematic figure of workflow in fMRelI. After defining study design and relevant contrasts, regressors can be split for cross-sectional analyses. Analyses are possible from a global to a local (voxelwise) level. Resulting reliability can be summarized based on an anatomical atlas (default: Harvard-Oxford brain atlas plus AAL cerebellum; see CONN) or predefined threshold can employed to identify regions surpassing a minimum of required reliability.

#### Preprocessing for within-session analysis: split-half estimation

Since longitudinal data is not available in every project and low reliability might also be caused by a substantial delay between repeated measures, we used a split-half estimation procedure to calculate cross-sectional reliability. The defined regressor of interest (offer onsets) and the corresponding parametric modulator (subjective value of offer) were randomly split into two parts while other regressors remained untouched (see *Figure 2*). Afterwards, first-level statistics were re-estimated. Thus, we could analyze the reliability of the task-induced activation within and between sessions. To the best of our knowledge, split-half reliability has not been used to assess reliability of fMRI yet, but was evaluated before in other brain imaging modalities such as electroencephalography (EEG) and magnetoencephalography (MEG; (Groppe, Makeig, & Kutas, 2009)).

**Figure 2:**
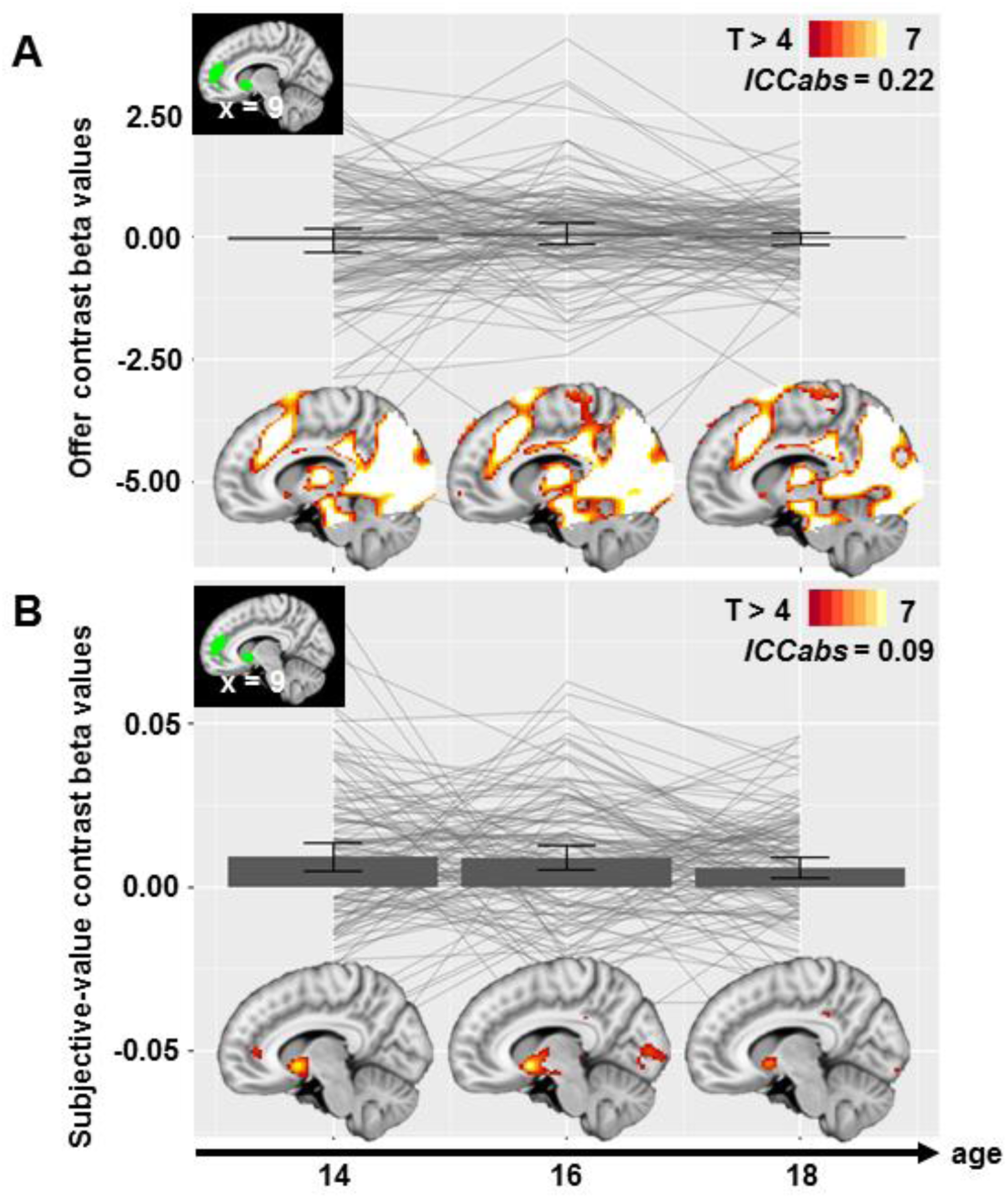
Individual trajectories of contrast (“beta”) values and the respective group activation maps for a region-of-interest (ROI, in green) in ventral striatum and vmPFC, derived from an independent sample using the same paradigm (Grosskopf et al., submitted), for the unmodulated (A) offer contrast and (B) the subjective value contrast. Nevertheless, signal derived from the offer contrast is more reliable in these regions. The ROI was based on an independent sample to avoid potential over-fitting and selection bias in reliability estimates.

#### Global reliability

A well-established approach to investigate fMRI reliability is the cluster overlap method. Here, a significance level is initially set to define “activated” voxels. Then, the degree of overlap in significant voxels between two measurements is quantified. We used the Dice and the Jaccard coefficient, which are commonly used in the literature. The latter can be easily interpreted as the percentage of overlapping significant voxels within all significant voxels (Jaccard, 1901; Maitra, 2010).

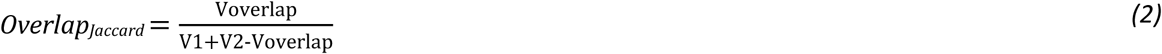

The Dice coefficient, first described by Rombouts and colleagues (1997), is defined as the number of voxels, which overlap (V_overlap_) divided by the average number of significant voxels across sessions (V_1_, V_2_).

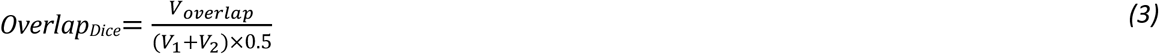

Both coefficients range from no overlap (0) to perfect overlap (1). To the best of our knowledge, there is no consensus for criterion indicating an “acceptable” overlap. In the review by Bennet and Miller (2010), the Dice coefficients ranged from 0.21 to 0.86 for various time lags between measurements (less than one hour up to 33 weeks), which may serve as a coarse reference for our results. Since the resulting reliability measures are strongly dependent on the significance threshold and the investigated data level, the toolbox offers the option to define the threshold and to calculate it at the individual or the group level.

Due to our specific interest in the difference between individual- and group-level data, we analyzed both levels using the uncorrected *p <* 0.01 as a rather liberal threshold to limit the initial loss of information due to thresholding.

As second global measure of reliability, we calculated the similarity of the fMRI activation maps, which was previously described as part of the representational similarity approach (Kriegeskorte, Mur, & Bandettini, 2008). Briefly, similarity between activation patterns has been used to characterize the resemblance of the neural representation of object and categories (Kriegeskorte et al., 2008). Likewise, Finn et al. (2015) adapted this approach to compare similarity of connectivity matrices across sessions to re-identify individuals based on maximum similarity (“fingerprinting”). They defined similarity as the Pearson correlation coefficient between vectors of edge values taken from the matrices of interest (Finn et al., 2015). In a similar manner, we compared brain activation matrices within and between subjects and sessions by vectorizing and correlating them. In other words, this procedure captures the resemblance of two patterns based on the alignment of high versus low brain activation values across the brain. Thus, we gathered information about reproducibility at the global level of the activation map. For each comparison in each contrast, we checked whether the within-subject similarity is the highest and therefore enables the re-identification of individuals. Since the similarities are correlation coefficients, they vary between a ‘perfect’ inverse relationship (-1.00) and a ‘perfect’ direct relationship (1.00). For statistical analyses, similarities were z-transformed.

#### Local reliability: ROI- and voxel-based measures

Besides the global approach, reliability can be estimated more granular for a specific region of interest (ROI) or for each voxel. In practice, the level of the reliability analysis should correspond to the level where content-related hypotheses are tested. In our case, we evaluated reliability within and outside the significant main effect regions because we expected that intra-individual stability would be to some extent independent of the magnitude of the group-level effect size. Therefore, we calculated each described measure at the voxel and ROI level. The ROI included ventral striatum and ventromedial prefrontal cortex (vmPFC) extracted based on an independent study using the same intertemporal choice paradigm (main effect of subjective value, *p*_*uncorrected*_ < .001, *k* > 20; Grosskopf, Kroemer, Böhme, & Smolka, submitted).

One important aspect of reliability is captured by the share of total variance, which is accounted for by the fact that repeated measures are nested within an individual or “class”. As introduced by Shrout and Fleiss (1979), the ICC is the coefficient of variance of interest and total variance. There are six forms of ICC differing in the defined variance of interest. We were primarily interested in the variance within participants and thus used two different types of ICCs, absolute (ICC_abs_) and consistency agreement (ICC_con_). The ICC_abs_ considers the session mean sum of squares (MS_session_) and therefore is the more conservative approach (see *Equation 4*), whereas the ICC_con_ is only relating the between-subject (MS_between_) to error (MS_error_) and within-subject mean sum of squares (MS_within_; see *Equation 5*). Here, k represents the number of sessions and n represents the number of subjects.

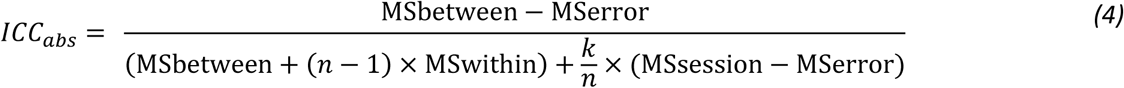

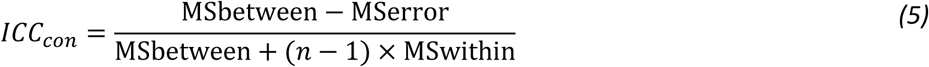

According to guidelines suggested by Fleiss (1986), ICCs lower than 0.4 represent poor reliability, ICCs between 0.4 and 0.75 represent fair (< 0.6) to good (>.6) reliability, and ICCs higher than 0.75 represent excellent reliability (Cicchetti, 2001).

In addition to ICCs, Pearson’s and Spearman’s correlation coefficient were calculated. Pearson’s and Spearman’s correlation coefficients are well-known measures for the strength of association between two variables. Pearson's correlation coefficient is the covariance of two variables, i.e. two vectors containing individual estimates in one voxel for two sessions, divided by the product of their standard deviations. Spearman’s correlation coefficient is defined by the Pearson’s correlation of the ranked variables. Thus, Pearson’s correlation represents the stability of the values (in interval scale) whereas Spearman’s correlation represents the stability of the rank order of the values (see *Equation 6*). The choice of the correlation coefficient as reliability marker is dependent on the assumptions and applications. If we expect a linear association, Pearson’s correlation is recommended. Moreover, Pearson’s r is often requested as input to conduct power analyses for repeated measures designs (e.g., in GPower; Faul, Erdfelder, Lang, & Buchner, 2007). Otherwise, Spearman’s correlation might be the coefficient of choice as it is more robust to non-linearity of changes across the range.

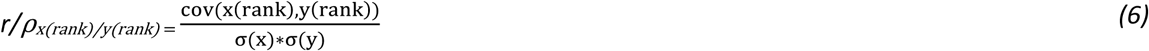

For the analysis of cross-sectional data, we applied the Spearman-Brown correction for split-half reliability (Spearman, 1910; see *Equation 7*), accounting for the underestimation of reliability due to the decreased number of items.

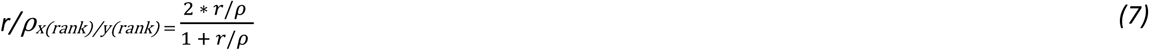

Here, we consider correlations (in absolute value) up to 0.35 as low, up to 0.67 as moderate and above 0.67 as high (Taylor, 1990). Despite their considerable descriptive value, systematic errors might lead to distortion of the coefficients. For example, if the second activity estimate was twice the first one, this would result in a decreased *r* (Müller & Büttner, 1994; Safrit, 1976). For general purposes, the ICC thus appears as more suitable.

#### Summary measures

To aid the assessment of local differences in reliability, several means of aggregation could provide useful insights. After calculating the voxel-wise correlations and ICCs, researchers may identify reliable regions by setting minimum reliability thresholds and a minimum cluster extent to restrict the following analysis to these regions narrowing down the set of candidate voxels (Fröhner et al., in prep).

In addition, recent research indicates that anatomical ROIs might produce more reliable data than ROIs generated out of functional data (Nord et al., n.d.). Hence, we created a reliability summary for all regions included in the atlas provided with CONN functional connectivity toolbox (Whitfield-Gabrieli & Nieto-Castanon, 2012) consisting of the Harvard-Oxford brain atlas (Desikan et al., 2006; Frazier et al., 2005; Goldstein et al., 2007; Makris et al., 2006) and the Automatic Anatomic Labeling (AAL) for the cerebellum (Tzourio-Mazoyer et al., 2002). Thus, the atlas-based summary might enable us to choose a sufficiently reliable anatomical ROI for future analyses and facilitates the comparison of reliability, for example between cortical and subcortical regions or across paradigms. Both, the identification of reliable clusters and the reliability summary for anatomical ROIs are implemented in fMRelI.

To visualize the summary, we grouped ROIs according to their maximum overlap with functional networks introduced by Yeo (Yeo et al., 2011). Furthermore, subcortical limbic regions were assigned to the limbic network and we included the cerebellum as an additional network. Finally, we correlated the split-half reliabilities of the offer and the subjective-value contrasts for each region to test if reliabilities at the ROI level are associated (i.e., statistically dependent).

#### Statistical dependence of reliability and signal amplitude and variance

Since we reasoned that effects at the group level are not necessarily predictive of the reliability at an individual level, we further assessed to what extent the local reliability is related to the average amplitude (“beta”) and its variance across individuals. Therefore, we correlated these group-level summary statistics with the z-transformed split-half reliability (*Spearman’s rho*) for each voxel. To emphasize the overall association, we show associations pooled across sessions.

## Results

To determine the reliability of a commonly employed intertemporal choice task across adolescence, we investigated the longitudinal trajectories of brain response to delayed monetary offers and the split-half reliability at each wave as a reference reflecting momentary reliability. The rationale behind this two-fold approach was to dissociate potential unreliability due to differential developmental trajectories from cross-sectional reliability of the measurement. We reasoned that retest reliability would not exceed cross-sectional reliability. Furthermore, to verify the scripts and constrain plausible results for the reliability of the longitudinal data, we simulated changes using known inputs of signal relative to varying degrees of noise (i.e., half to double the initial signal-to-noise ratio).

### Main effect results for subjective value and offer contrast

First, we observed a strong main effect of the offer onset event, which was similar across the three sessions, encompassing the occipital and parietal cortex, the thalamus, and the dorsal anterior cingulate cortex (dACC). Second, we observed that the parametric modulator “subjective value” of the offer event was associated with activation in ventral striatum (peak: −9, 5, −6; *T*_*max*_ = 11.02; *p*_*FWE-corrected*_ < 0.05), vmPFC (peak: −3, 41, 4; *T*_*max*_ = 9.28; *p*_*FWE-corrected*_ < .05) and posterior cingulate cortex (peak: 0, −31, 34; *T*_*max*_ = 7.52; *p*_F*WE-corrected*_ < 0.05). Hence, in line with numerous previous studies (e.g., Koffarnus et al., 2017; McClure, Laibson, Loewenstein, & Cohen, 2004; Peters & Buchel, 2009), there were robust and seemingly congruent main effects for both contrasts across sessions *(Figure 2)*.

### Global reliability

To assess the overlap of activation maps across sessions, we calculated the Jaccard and Dice coefficients as indices of similarity a) individually for each subject and b) aggregated at the group-level at a liberal threshold (*p*_*uncorrected*_ < .01; see *Table 1*). For the offer contrast, the Dice overlap ranged from .91 to .96 at the group level, whereas it ranged from .39 to .61 for the subjective-value contrast. Critically, the congruency of group main effects was in stark contrast to the unreliability of individual activation maps. In particular, the parametric contrast for subjective value yielded very low coefficients of overlap, suggesting that the individual information contained in the contrasts is not reliable (*Figure 3*). Applying more stringent thresholds also lead to worse overlap coefficients (not reported), indicating that these results are not due to a liberal threshold at the individual level. The notable difference between individual and group-level stability was also visible in the cross-sectional comparison alone substantiating that the low reliability is not simply explained by low stability of the contrast estimates across time during adolescence. Notably, the low reliability of the brain response associated with parametric value tracking stands in contrast to the behavioral results, where the ICCs for the discount rate were well within the moderate range (decisions during pretest: 0.57, decisions during fMRI: 0.44).

**Table 1:** Comparison of average individual overlap and group-level overlap for offer and subjective-value contrast

| Comparison |  | Group level |  | Mean individual level |  |
| --- | --- | --- | --- | --- | --- |
|  |  | Jaccard* | Dice* | Jaccard* | Dice* |
| Offer contrast |  |  |  |  |  |
| Longitudinal | 14 to 16 | 0.85 | 0.92 | 0.17 | 0.26 |
|  | 14 to 18 | 0.83 | 0.91 | 0.17 | 0.25 |
|  | 16 to 18 | 0.87 | 0.93 | 0.18 | 0.27 |
| Cross-sectional | Split 14 | 0.89 | 0.94 | 0.12 | 0.17 |
|  | Split 16 | 0.91 | 0.96 | 0.13 | 0.18 |
|  | Split 18 | 0.90 | 0.95 | 0.10 | 0.15 |
| Subjective-value contrast |  |  |  |  |  |
| Longitudinal | 14 to 16 | 0.37 | 0.54 | 0.02 | 0.02 |
|  | 14 to 18 | 0.30 | 0.46 | 0.01 | 0.02 |
|  | 16 to 18 | 0.35 | 0.52 | 0.01 | 0.03 |
| Cross-sectional | Split 14 | 0.31 | 0.47 | 0.01 | 0.02 |
|  | Split 16 | 0.44 | 0.61 | 0.02 | 0.03 |
|  | Split 18 | 0.56 | 0.39 | 0.01 | 0.02 |

Note. \*Overlap coefficients are calculated for suprathreshold voxels ( $p_{uncorrected} < .01$ ).

**Figure 3:**
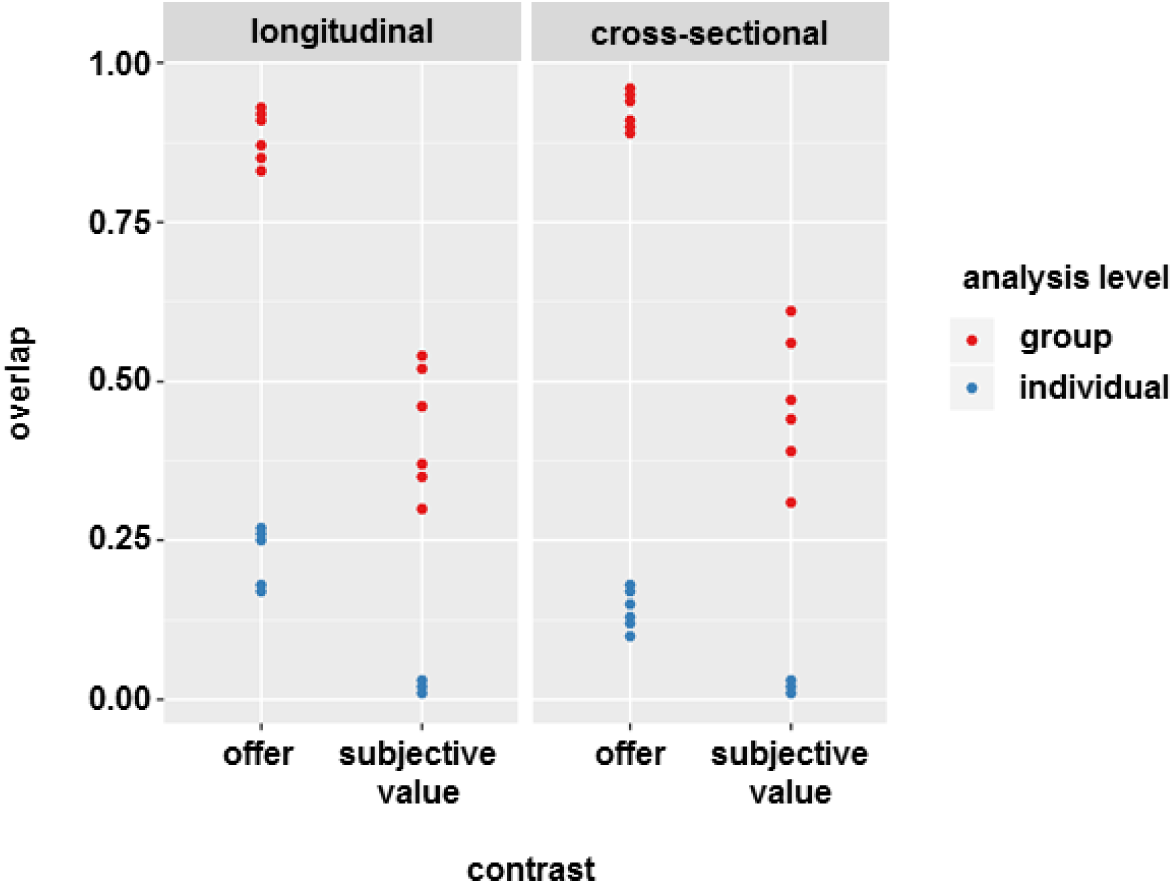
Dot plot of overlap coefficients separated for longitudinal and cross-sectional analysis, group and individual and offer and subjective value contrast. Overlap is lower for the subjective-value contrast compared to the offer contrast and lower at the individual level compared to the group level.

Due to the marked difference between reliability at the group vs. individual level, we next calculated the similarity between individual activation maps across individuals and sessions (z-transformed for statistical analyses). Conceptually, this method enabled us to quantify the resemblance of individual brain activation patterns across individuals and contrasts demonstrating how unique an induced brain response is. Statistically, this method enabled us to quantify within- and between-subject similarity of task-evoked brain activation without the necessity to define an arbitrary statistical threshold. Again, we observed higher similarity within subjects (.60 ≤ r_average_ ≤ .66) compared to other subjects (.18 ≤ r_average_ ≤ .20) for the offer contrast map, *T(125)* ≥ 17.6, *p* < .001 (*Figure 4*). Although the difference between similarity within subjects (.07 ≤ r_average_ ≤ .08) compared to other subjects (r_average_ ≈ .01) was also significant for the subjective-value contrast, *T(125)* ≥ 2.1, *p* < .037, within-subject similarity was significantly higher for the offer contrast, *T(125)* ≥ 10.0, *p* < .001. Similar results were obtained for cross-sectional data using the split-half method (*Figure 4*; see *Table S.2* and *S.3* for complete T-test results).

**Figure 4:**
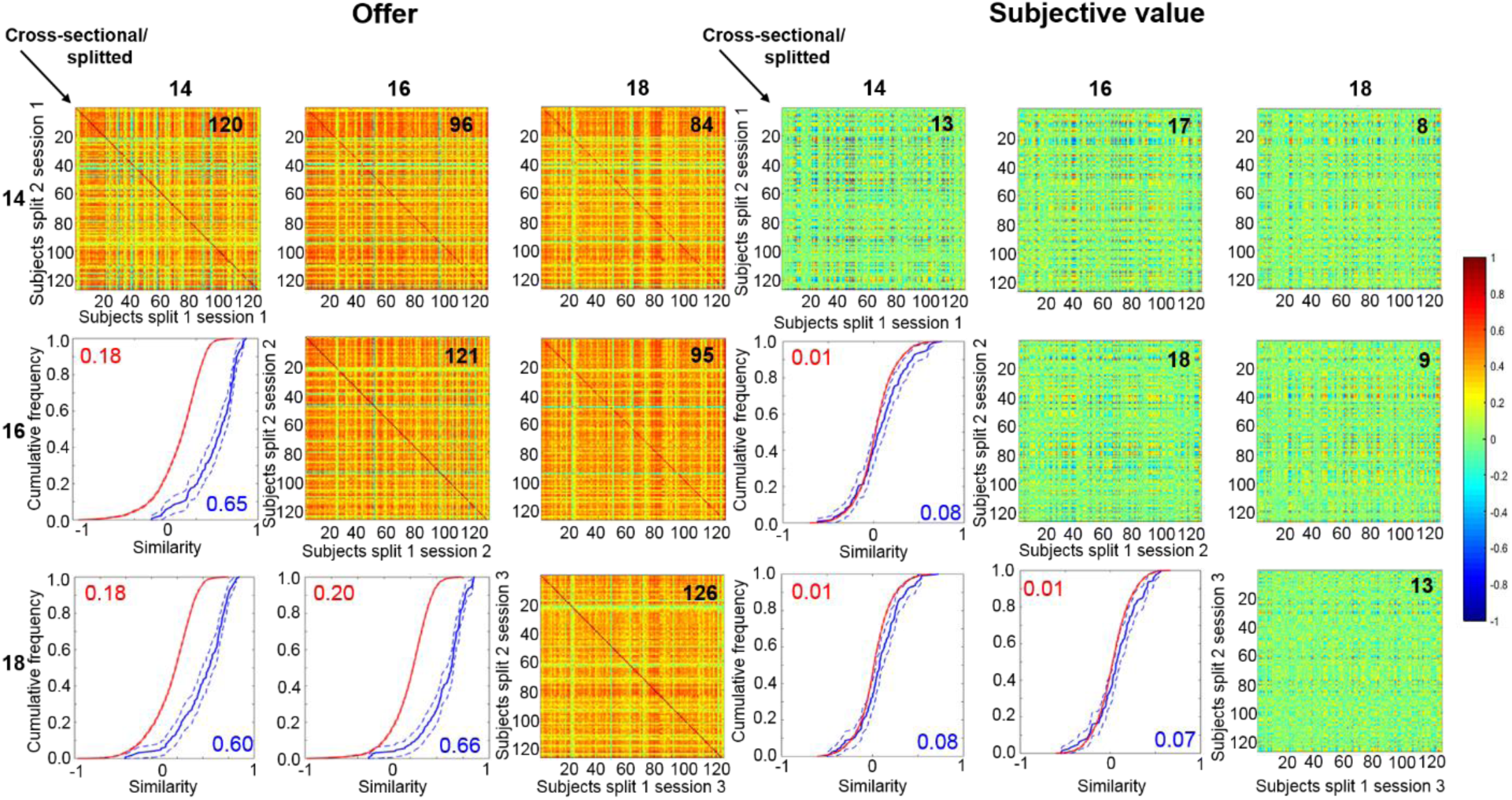
Similarity maps for all longitudinal and cross-sectional comparisons. Each row represents the activation map of one subject for one session or half of a session (split). Each column represents the activation map of one subject for another session or the second half of the session. For reliable activation “fingerprints”, we would expect the highest similarity estimates for the within-subject comparison. This would be visible in a prominent diagonal as is the case for the offer contrast (left panel). This is also evident in the empirical cumulative density functions (ecdf; lower diagonal): If the density of within-subject similarity (blue) differs from between-subject similarity (red), we should see an offset between the two lines. The higher the similarity is on average, the more the lines will be shifted to the right. The last column represents the similarity to the respective mean image. The numbers on the matrices indicate the number of correctly re-identified subjects, which is considerably higher for the offer contrast.

To further disentangle sources of variability, variance analyses with the factors level (within and between) and contrast (subjective value and offer) revealed a significant interaction between level and contrast for all comparisons, *F* ≥ 147.9, *p* < .001 (see *Figure 5* and supporting information *Table S.1* for complete ANOVA results). In general, similarities were lower for the subjective-value contrast than for the offer contrast, *F* ≥ 659.7, *p* < .001. Thus, the difference between within- and between-subject similarities is higher for the offer contrast, which indicates that the signal elicited during the offer contrast contains more unique individual information compared to the subjective-value contrast.

**Figure 5:**
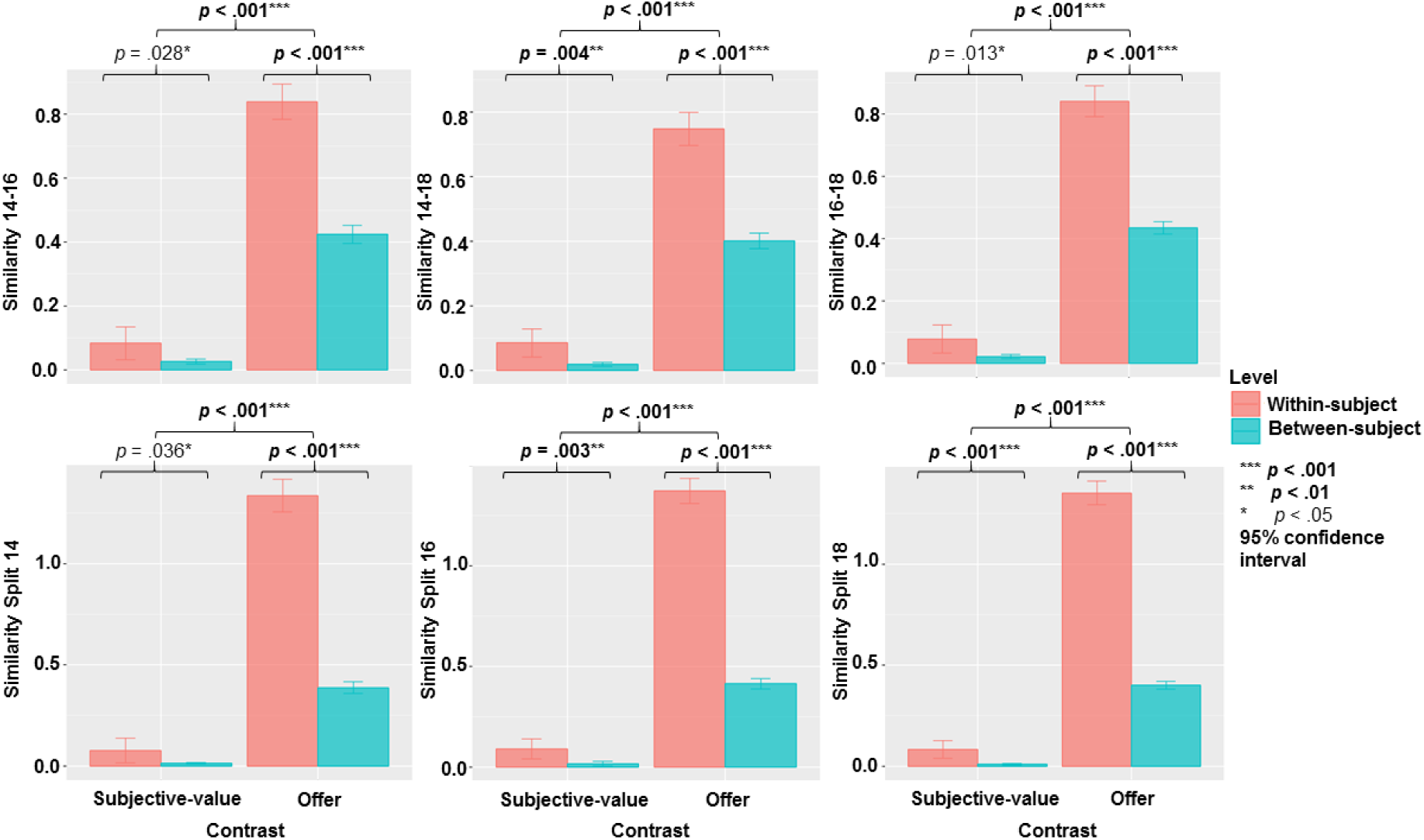
Summary of z-transformed similarities for each comparison: stars represent significance of paired T-tests for within- and between-subject similarities (offer: *T* ≥ 17.6, *p* ≤ .001; subjective value: *T* ≥ 2.1, *p* ≤ .036) and for the difference between within- and between-subject similarities, which were higher for the offer contrast than for the subjective-value contrast (*T* ≥ 10.0, *p* ≤ .001; see supporting information)

In the next step, we examined whether we could re-identify individuals based on maximum similarity. Again, there was a clear difference between the offer contrast and the subjective-value contrast. For the offer contrast, 84 to 126 (i.e., all) subjects (ranging across comparisons) could be re-identified based on their maximum similarity to a second scan, whereas it was only the case for 8 to 18 subjects for the subjective-value contrast (see *Figure 4*). Note that according to a binomial distribution, ≥3 correct classifications would be considered as better than chance (*p* < .0185) indicating that both contrast work significantly better than chance in re-identifying individuals.

In addition, we assessed similarities for the motor contrasts, which yielded very similar results to the offer contrast (see supporting information, *Figure S.1*) that is substantially higher within-subject compared to between-subject similarity. Lastly, we ran simulations with known signal (i.e., individual activation maps at age 14), which were corrupted by noise mimicking changes over time primarily due to measurement error. Even for the highest level of noise added to the individual contrast maps (i.e., double the initial variance), within-subject reliability was preserved to a moderate extent suggesting that the absence of within-subject reliability is more than a simple result of noisy test-retest data (*Figure 6*).

**Figure 6:**
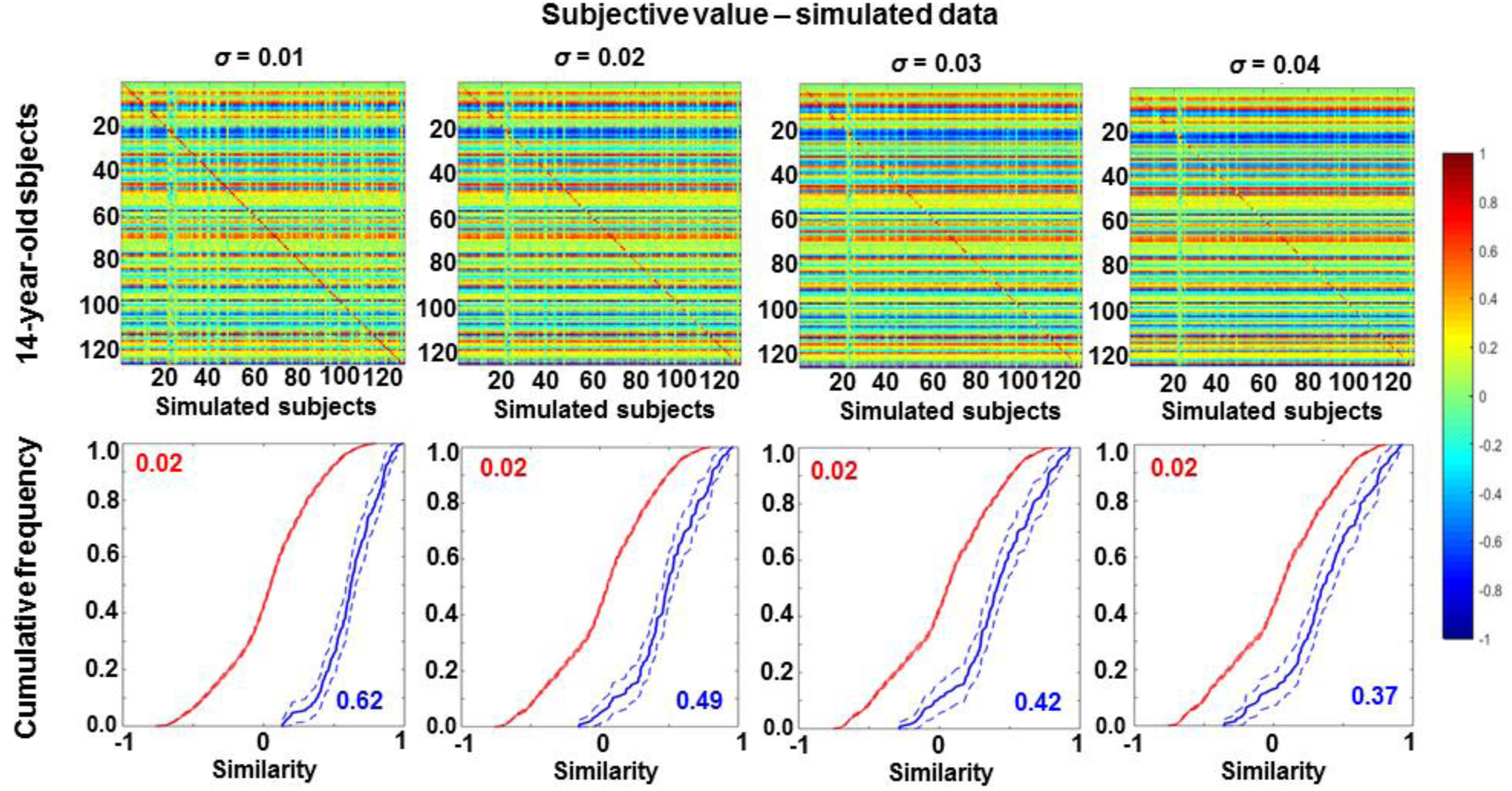
Similarity maps and empirical cumulative distribution functions (ecdfs) for the comparison of the 14-year-old and simulated changes in the subjective-value contrast with increasing noise from *σ* = 0.01 (half the initial variance) on the left to *σ* = 0.04 (double the initial variance) on the right. Whereas increasing levels of noise in the individual estimates lead to less within-subject similarity (diagonal and blue lines), excessive measurement noise alone cannot explain the low reliability that we observed in the subjective-value contrast.

To conclude, the task elicits reliable brain activation patterns, yet the subjective-value contrast fails to achieve the required minimum level of reliability. This suggests that the low reliability is not due to the collected data, the task, or high measurement noise *per se*, but rather attributable to the specific parametric contrast supposed to track subjective value.

### Local reliability

Next, we sought to identify regional differences in the reliability of task-evoked brain responses by calculating ICCs and correlations for each voxel. Across the brain, ICCs and correlations were much lower for the subjective-value contrast than the offer contrast and results were highly similar for the different longitudinal and cross-sectional correlation coefficients (*Figure 7*). Regional differences in reliability were visible for the offer contrast, where we observed higher correlations in visual and parietal regions and lower correlations in orbitofrontal regions. Notably, there was no such apparent pattern of regional differences in the reliability for the subject-value contrast. Even in the commonly identified value-tracking regions, we find higher average correlations for the offer contrast compared to the subjective-value contrast.

**Figure 7:**
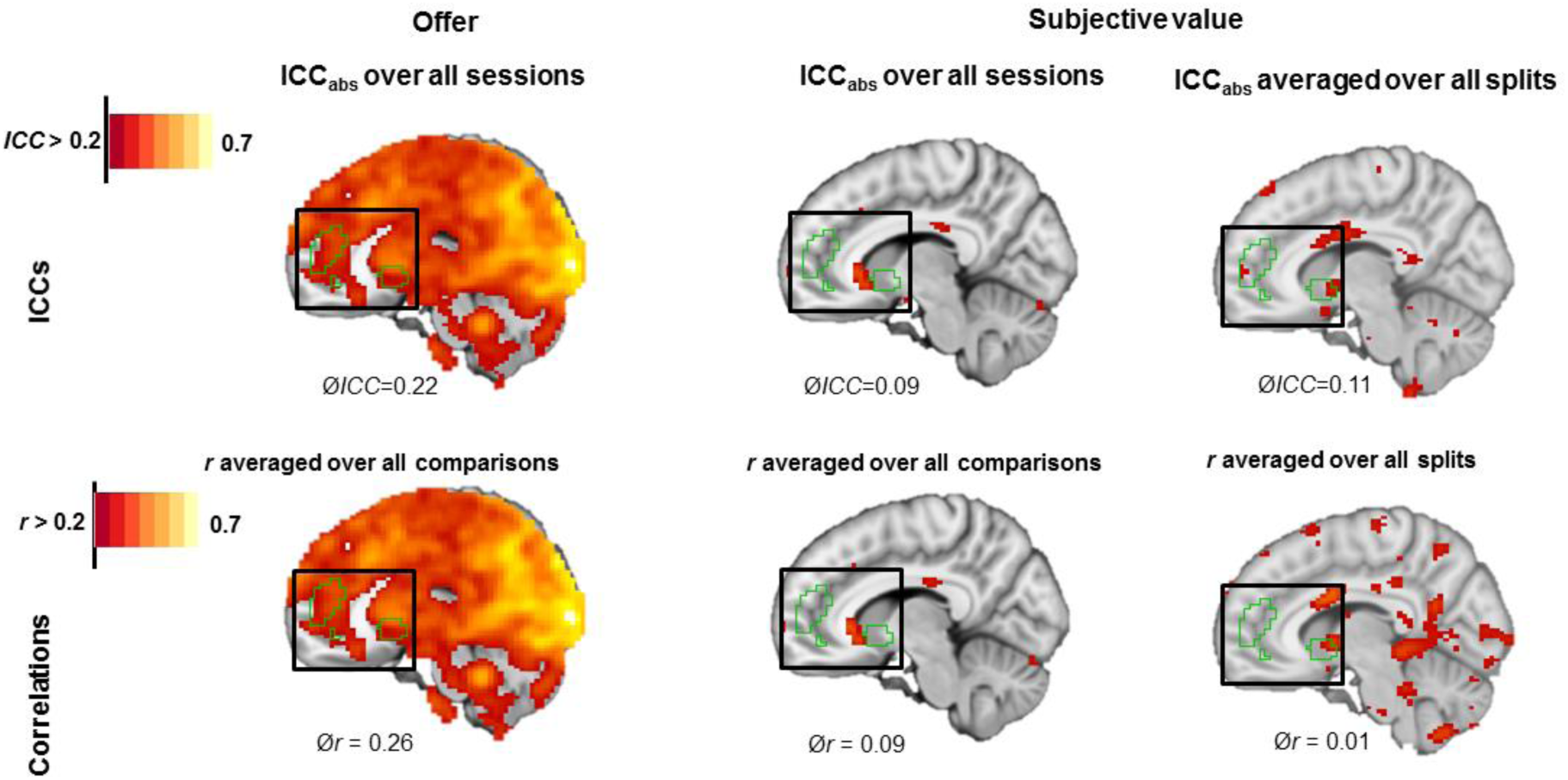
*ICC*_*abs*_ and Pearson’s *r* over all sessions (longitudinal) for both contrasts and for the averaged split (cross-sectional) data for the subjective-value contrast depicted for selected sagittal slice. Average values indicate reliability within the independently identified ROI encompassing ventral striatum and vmPFC (green outline).

To summarize all voxel-wise data, we created a matrix with an average ICC and correlation for each region included in the atlas. For visualization purposes, we grouped the ROIs in functional networks according to Yeo (Yeo et al., 2011). Again, we could see that the reliability in the offer contrast was higher compared to the subjective-value contrast (see *Figure 8*). Moreover, network-based differences in reliability occurred for the offer contrast (highest in the visual network), but there was no indication of network-based differences in reliability for the subjective-value contrast. The correlation of split-half reliabilities of both contrasts did reveal a significant, but weak positive association (*r* = .14, *p* < .001; see *Figure 8*). Notably, reliability only surpassed the moderate criterion for both contrasts in the nucleus accumbens in 18-year-old participants.

**Figure 8:**
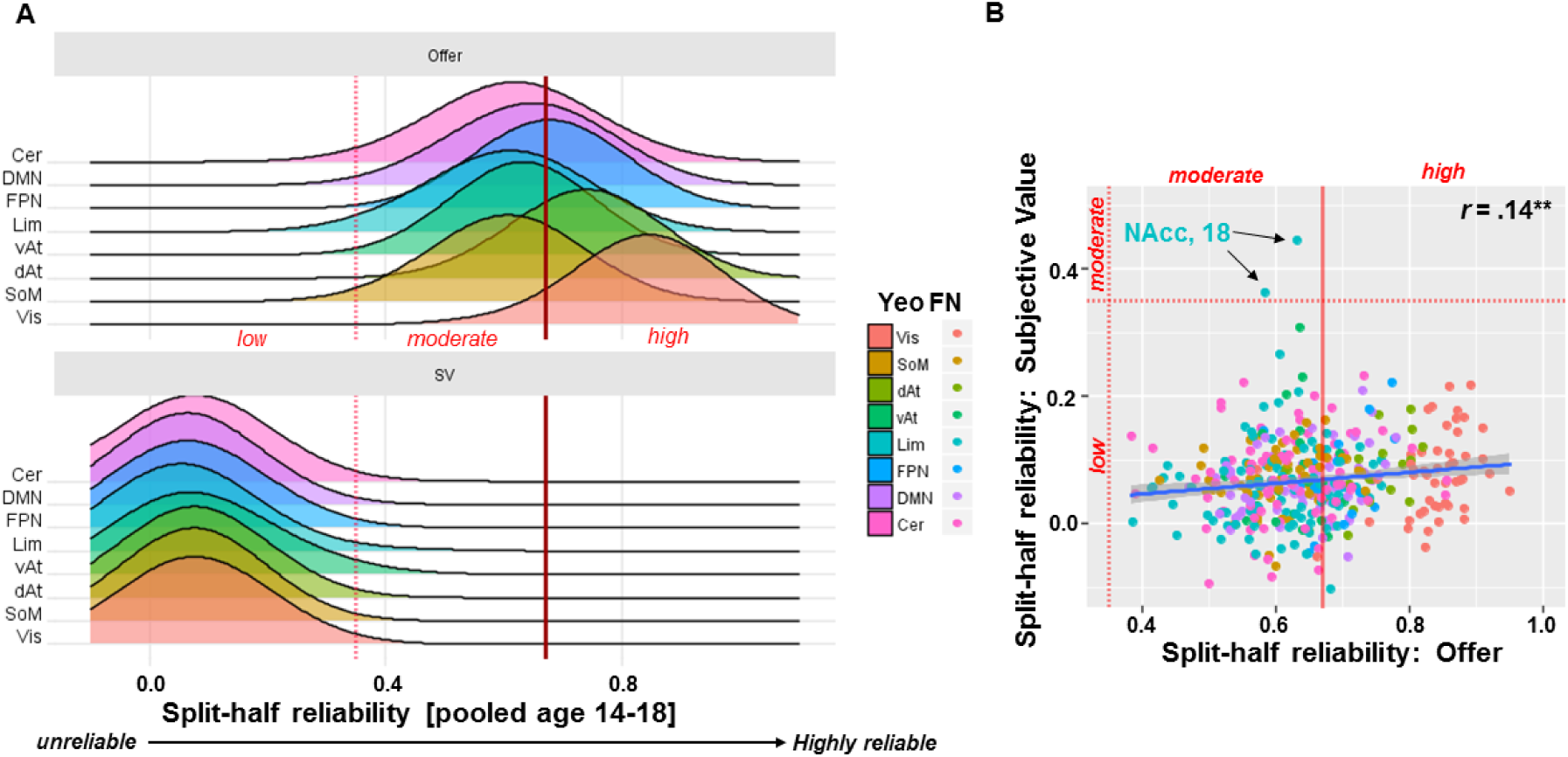
(A) Density plots of the split-half reliabilities (*r*) for offer and subjective-value (SV) contrasts over all assessments. (B) Correlation between split-half reliabilities (*r*) for offer and subjective-value contrasts in each region listed in the atlas. Reliability for both contrasts only exceeded the moderate criterion within the nucleus accumbens (NAcc) of 18-year-old participants. (A+B) colored according to Yeo’s (2011) functional networks (FN): visual (Vis), somatomotor (SoM), dorsal attention (dAt), ventral attention (vAt), limbic (lim; including subcortical limbic regions), frontoparietal (FPN), default mode (DMN), cerebellum (Cer; added as network).

### Statistical dependence of reliability and signal amplitude and variance

A common assumption is that greater amplitude of brain response is associated with higher reliability. Yet, empirical evidence for this assumption is scarce and our results indicate a disconnection between group-level congruence and individual reliability. To further elucidate the role of signal amplitude and variance for the reliability analysis, we correlated the split-half reliability (*r*) of each session with the average amplitude and variance of the signal across subjects within each gray-matter voxel.

In line with the expected association, we found that higher average betas were associated with greater split-half reliability for the offer contrast (Spearman’s rho = .55), but only to a negligible extent for the subjective-value contrast (Spearman’s rho = .06). Further in line with our hypothesis that greater inter-individual variability is important for reliability at the individual level, we observed a positive association between the two for the offer contrast (rho = .57) and, to a weaker extent, for the subjective-value contrast as well (rho = .18). Perhaps surprising at first, this leads to an attenuated rank-order correlation for t-values with reliability (offer: rho = .45, subjective value: rho = .00; see *Figure 9*) compared to average amplitude or inter-individual variability alone. An additional exploratory analysis where we orthogonalized inter-individual variability and amplitude using linear regression indicated that higher reliability was observed in voxel with lower or higher inter-individual variability in brain response than what would be expected from a linear association (see *Figure 10*). Taken together, these results may indicate that a substantial degree of the reliability in contrasts that show good psychometric characteristics can be accounted for by signaling characteristics.

**Figure 9:**
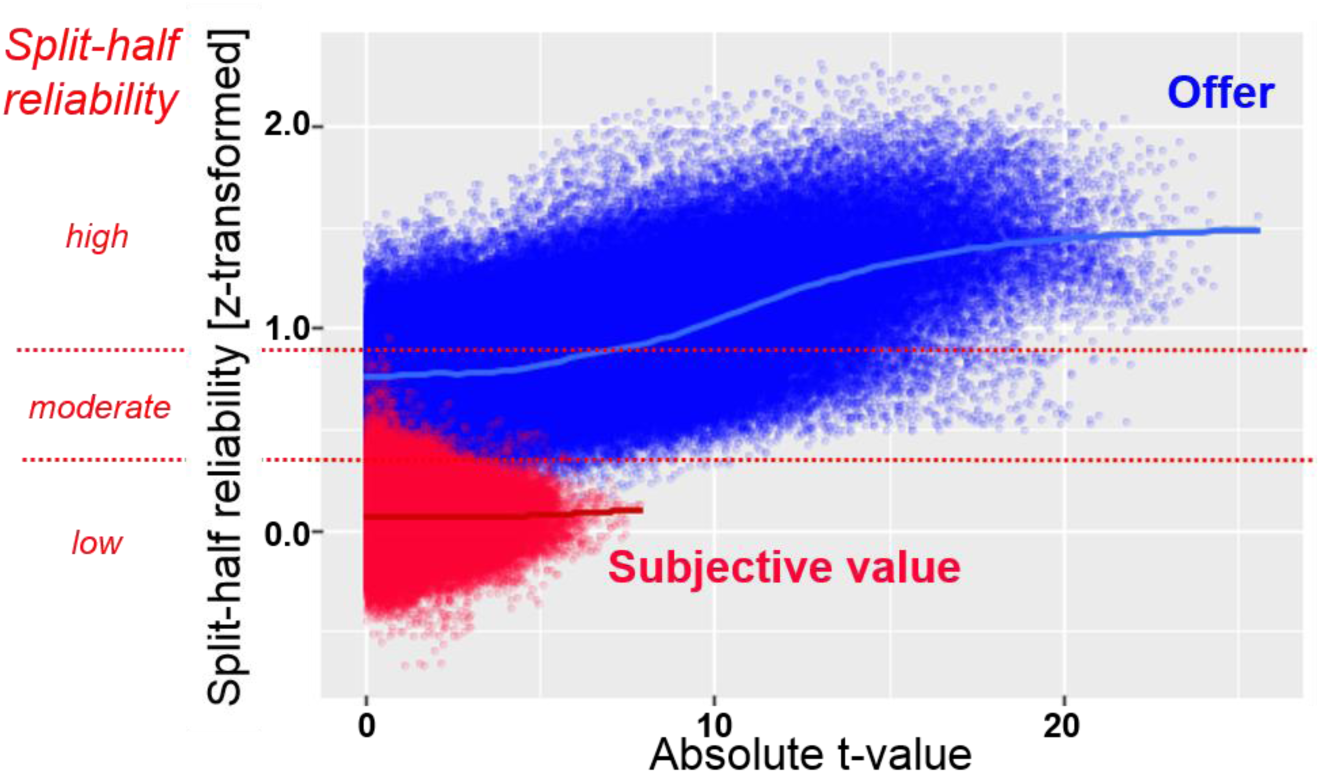
Association of t-value and split-half reliability for offer (blue) and subjective value (red). Note that for t-values of comparable magnitude, voxelwise reliability is lower in the subjective-value contrast compared to the offer contrast.

**Figure 10:**
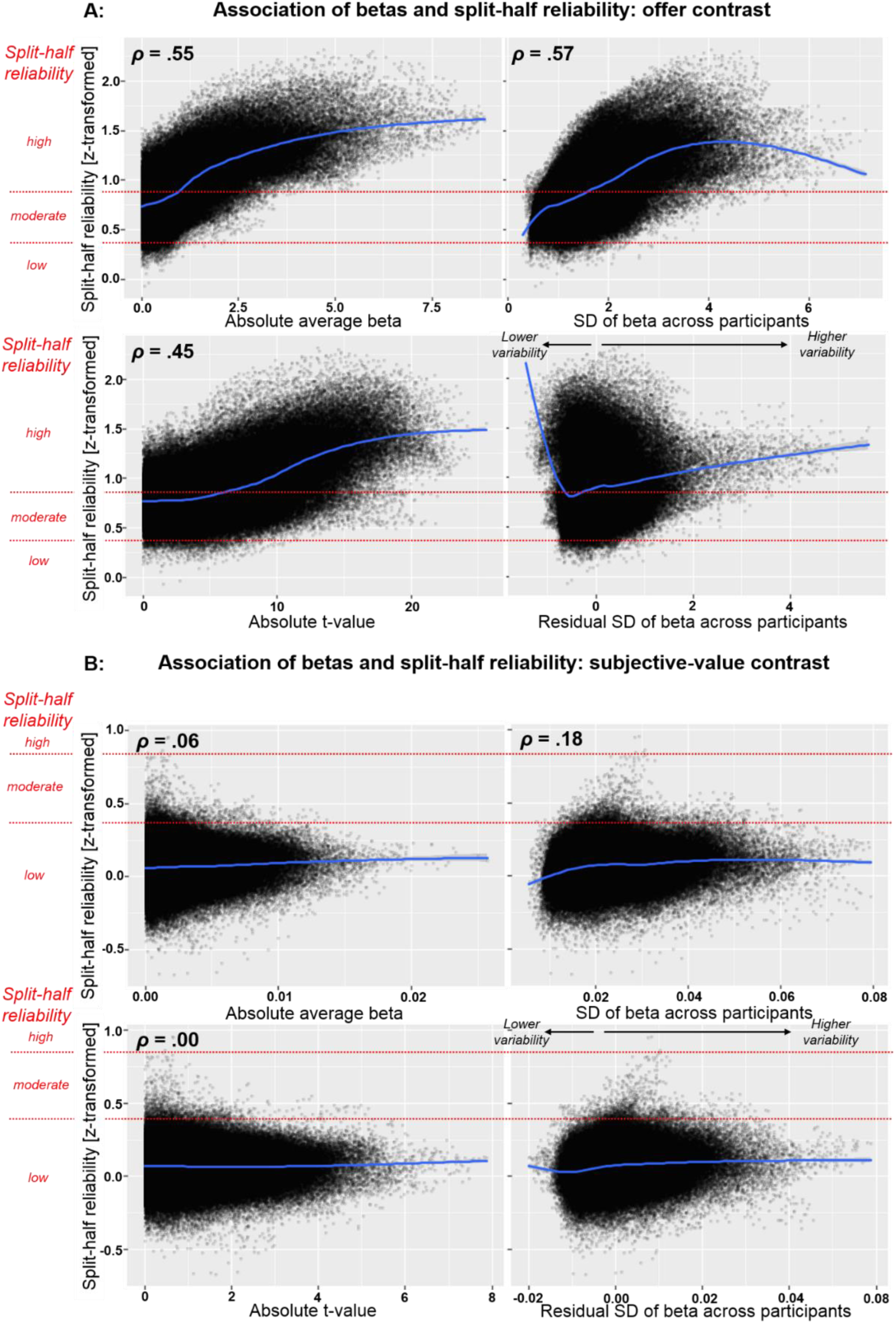
Moderately strong associations between signal amplitude (beta and t-value), inter-individual variability (SD and residual SD) and split-half reliability for the offer contrast (A), but only weak associations for the subjective value contrast (B).

## Discussion

Reliability is a key aspect of the diagnostic quality of any measurement such as a biomarker derived from brain activation to monetary offers. Yet, despite the recent surge of interest in biomarkers, little is known about the reliability of many common paradigms, which are frequently used in fMRI research. Here, we provided a comprehensive approach to investigate cross-sectional (within-session) and longitudinal (between-session) reliability of fMRI data. Using longitudinal data of an intertemporal choice task, we described an extensive analysis of reliability spanning from the group to the individual level and from the global to the local level. The key results were consistent across the applied measures: First, the individual reliability of the brain response to the offer onset is substantially higher than the reliability of the parametric value-tracking signal. Second, group-level reliability is substantially higher than individual reliability. Third, cross-sectional and longitudinal reliability of the subjective-value contrast were similarly low and failed to achieve a minimum level of reliability that is required for use as a biomarker. Fourth, we provide preliminary evidence that reliability is positively dependent on signal amplitude and inter-individual variability emphasizing the necessity to optimize between-subject variance for reliable classification and prediction.

### Good reliability at the group level versus unreliability at the individual level

In line with previous studies on the stability of group effects (Bennett & Miller, 2010; Freyer et al., 2009; Nord, Gray, Charpentier, Robinson, & Roiser, 2017; Plichta et al., 2012), we observed congruent brain activation for both onset and subjective value of the delayed offer over time. However, individual trajectories were not reliable over time as well as inconsistent across split halves of each session. The difference between reliability at the individual versus the group level was even more pronounced in the subjective-value contrast. Since our study is not the first revealing such notable difference between individual- and group-level reliability (van den Bulk et al., 2013; Vetter et al., 2015), these results call for caution in using group-level results to draw conclusions about individual aspects of brain function. Illustratively, increased group-level brain activity in a specific brain region during the choice of one option does not necessarily imply that these choice “signatures” would suffice to predict choice preferences at the individual level in terms of reliability. Thus, our results might explain the limited success in connecting neural and behavioral results for many commonly employed tasks in cognitive neuroscience (e.g., Müller et al., 2015; Nebe et al., 2017; Whelan et al., 2014), perhaps due to an emphasis on paradigms producing robust group effects.

Whereas differences between the individual versus the group level were striking, multiple sources are likely to contribute to the observed pattern. For example, greater measurement noise contained in the individual compared to the averaged group-level data could be essential. However, in our simulation, we doubled the inter-individual variability of the first-wave estimates and were still able to recover a considerably more reliable rank order compared to the stability of the subjective-value contrast. Moreover, good reliability and whole-brain similarity as evident in the offer or motor contrasts speaks against a simple measurement noise account. Hence, the simulation and the data from simple contrasts does not support the idea that fMRI data in general is too noisy to be used as a biomarker for individualized prediction in clinical research. Nevertheless, it emphasizes that not all contrasts are equally useful at an individual level, at least according to their basic psychometric characteristics. Similarly, our results demonstrate that not only the amplitude of the brain signal but also the inter-individual variance in the signal is positively associated with the reliability. Hence, inter-individual variance is needed to capture brain activation as an individual characteristic and to differentiate it from a commonly shared neural evaluation process.

### Use of reliable brain activation patterns as a potential tool for clinical research

In the search for potential biomarkers of mental disorders, reliability at an individual level is a major limitation in gauging its potential predictive value. Hence, we should expect a notably higher within-subject compared to between-subject reliability for any promising candidate such as the value-tracking signal (Ripke et al., 2014). In our case, the within-subject similarity was indeed higher than the between-subject similarity. However, the absolute values and the difference between individual and group levels was much lower for the subjective-value contrast and largely failed to exceed the recommended minimum threshold for moderate reliability. Whereas developmental changes could partly explain low longitudinal reliability occurring between sessions, such changes cannot explain the low consistency across split halves of the paradigm within a given session. Also, the high intra-individual similarity and correspondingly high accuracy in re-identifying participants using brain activation patterns derived from the offer contrast suggests that there is a distinct individual component contained in the processing of monetary offers. Intriguingly, the congruent group activation in the subjective-value contrast might indicate that the underlying value-tracking signal commonly seen in the ventral striatum and vmPFC could be immutably shared among participants. This would explain why an unreliable individual response occurs within a well-replicated network consistently reproduced at the group level, which is also conclusively supported by other neuroscientific methods including decades of animal research (e.g., Floresco, 2013; Floresco, Maric, & Ghods-Sharifi, 2008; Hamid et al., 2016; Saddoris et al., 2015). Taken together, these results suggest that the parametric contrast for subjective value is not a suitable candidate as a biomarker for individualized prediction because it fails basic diagnostic criteria, whereas the offer contrast achieved a sufficiently high reliability across individuals and sessions.

More generally speaking, a weak correspondence between the estimated brain response in two independent halves, runs, or sessions indicates a failure in reliably differentiating between individuals. In turn, this makes it improbable to detect associations with other more distant outcomes (Hedge et al., 2017). During the last decade, several large-scale studies started to investigate the predictive validity of fMRI data for select aspects of human behavior such as the IMAGEN project (Schumann et al., 2010), the UK biobank (www.ukbiobank.ac.uk; Sudlow et al., 2015) or the Human Connectome Project (Van Essen et al., 2012). Whereas the predictive validity of many classic paradigm may turn out to be limited, these large studies offer the possibility to investigate such paradigms systematically regarding psychometric characteristics, which will ultimately help in identifying paradigms, contrasts, and conditions yielding a sufficiently high reliability. To this end, our toolbox fMRelI may facilitate the development of a standardized and comprehensive approach in establishing the reliability of biomarkers derived from fMRI data.

### Limitations

The current study was limited to one exemplary paradigm. Thus, the results may not generalize to other tasks. Still, the intertemporal choice paradigm provides a representative case as it is often used in fMRI research and we had sufficient data across three waves of data collection (i.e., >1h of task-based fMRI data per participant). The high consistency of the obtained reliability estimates across waves further corroborates the evidence provided by this key example for other relevant scenarios. Thus, we feel that our results warrant to call for more caution in future research targeting individual aspects of brain function, particularly in longitudinal studies. Further investigations may contribute to specific aspects of the design that determine individual reliability of more nuanced facets of value tracking. Second, new methods for the estimation of fMRI reliability have been proposed that may provide additional insights (Maitra, Roys, & Gullapalli, 2002; Shou et al., 2013; Zandbelt et al., 2008). So far, we primarily collected the most commonly employed reliability indices in a new toolbox to make them more readily accessible for fMRI research, but other methods should be implemented in the future that might make better use of the unique characteristics of fMRI data. To this end, we welcome the addition of novel measures to the toolbox to accelerate the dissemination among the fMRI community. Third, other statistical methods incorporating hierarchical priors on parameter distributions that improve the recovery of brain response estimates could be employed in the future to improve the accuracy of individual estimates (e.g., Kroemer et al., 2014, 2016).

## Conclusions

Using an intertemporal choice task, we have shown that there is a substantial difference between group-level and individual-level reliability in brain response to monetary offers and that the extent of that difference varies strongly between contrasts within the task. Simple contrast reflecting activation elicited by the presentation of monetary offers or motor responses showed good reliability and allowed us to re-identify individuals with high accuracy across multiple waves. Critically, the subjective-value contrast, which is commonly used to assess individual differences in value tracking showed insufficiently low reliability across multiple indicators and levels of analysis. To conclude, our results suggest that promising biomarkers should be extensively evaluated with respect to intra-individual stability over time before they can be routinely applied for prediction or classification to avoid the reliability fallacy arising from congruent group activation maps. Importantly, we provide all the functions that we have employed in the MATLAB-based toolbox fMRelI to facilitate the use in future analyses of fMRI reliability across many more applications in cognitive neuroscience.

## Acknowledgements

This study was supported by the Deutsche Forschungsgemeinschaft (DFG Grant # SFB 940/1 & SFB 940/2 and the German Ministry of Education and Research (BMBF Grant # 01EV0711 & # 01EE1406B). JHF received a PhD-scholarship from the SFB 940 „Volition and Cognitive Control: mechanisms, modulators and dysfunctions”. VT and NBK were supported by the University of Tübingen’s fortüne program, grant #2453-0-0.

We thank Marie Stolze and Caroline Burrasch, who contributed to the implementation and documentation of the toolbox. We thank Stephan Ripke for his previous work on the presented intertemporal choice task.

## Supporting information

### Similarity output

In total, the toolbox offers three figures to illustrate similarity. The first figure is a color matrix, which represents the similarity estimates between and within each subject for each comparison. To aid identifying outliers, we included the similarity to the respective mean group image in the last column. The second figure is a histogram of within and between subject similarities (in correlation coefficients). The third figure is an empirical cumulative density function, which facilitates the identification where along the axis differences in similarity between subjects or contrasts arises. Accordingly, fMRelI creates the visual outputs automatically after computing the global similarities.

**Table S.1:** results of ANOVAs to identify differences in similarities between contrasts (offer and subjective value) and analysis levels (within- and between-subject)

| Comparison | Effect | *df* | *F* | *p* |
| --- | --- | --- | --- | --- |
| 14 to 16 | contrast | 1,125 | 914.8 | < .001 |
|  | level | 1,125 | 147.2 | < .001 |
|  | contrast x level | 1,125 | 184.8 | < .001 |
| 14 to 18 | contrast | 1,125.061 | 659.7 | < .001 |
|  | level | 1,125.110 | 185.0 | < .001 |
|  | contrast x level | 1,124 | 94.3 | < .001 |
| 16 to 18 | contrast | 1,125 | 1090.3 | < .001 |
|  | level | 1,125 | 213.4 | < .001 |
|  | contrast x level | 1,125 | 147.9 | < .001 |
| Split 14 | contrast | 1,125.071 | 873.5 | < .001 |
|  | level | 1,125.093 | 439.3 | < .001 |
|  | contrast x level | 1,124 | 432.3 | < .001 |
| Split 16 | contrast | 1,125 | 1480.7 | < .001 |
|  | level | 1,125 | 637.6 | < .001 |
|  | contrast x level | 1,125 | 685.7 | < .001 |
| Split 18 | contrast | 1,125.081 | 1934.3 | < .001 |
|  | level | 1,125.085 | 785.5 | < .001 |
|  | contrast x level | 1,124 | 790.3 | < .001 |

**Table S.2:** results of T-tests und identify differences between within- and between-subject similarities for the different comparisons and contrasts (offer and subjective value)

| Comparison | Effect | *df* | *T* | *p* |
| --- | --- | --- | --- | --- |
| 14 to 16 | offer | 125 | 19.8 | < .001 |
|  | subjective value | 125 | 2.2 | .028 |
| 14 to 18 | offer | 125 | 17.6 | < .001 |
|  | subjective value | 125 | 3.0 | .004 |
| 16 to 18 | offer | 125 | 20.3 | < .001 |
|  | subjective value | 125 | 2.5 | .013 |
| Split 14 | offer | 125 | 28.4 | < .001 |
|  | subjective value | 125 | 2.1 | .036 |
| Split 16 | offer | 125 | 34.3 | < .001 |
|  | subjective value | 125 | 3.0 | .003 |
| Split 18 | offer | 125 | 37.2 | < .001 |
|  | subjective value | 125 | 3.3 | < .001 |

**Table S.3:** results of T-tests und identify differences between contrasts (offer – subjective value) regarding difference of within- and between-subject similarities

| Comparison | *df* | *T* | *p* |
| --- | --- | --- | --- |
| 14 to 16 | 125 | 13.6 | < .001 |
| 14 to 18 | 125 | 10.0 | < .001 |
| 16 to 18 | 125 | 11.3 | < .001 |
| Split 14 | 125 | 20.8 | < .001 |
| Split 16 | 125 | 26.2 | < .001 |
| Split 18 | 125 | 28.1 | < .001 |

### Summary measures

Users have the opportunity to set their own thresholds for cluster size and reliability measure. Furthermore, one could replace the atlas.

#### Results motor contrasts

**Figure S.1:**
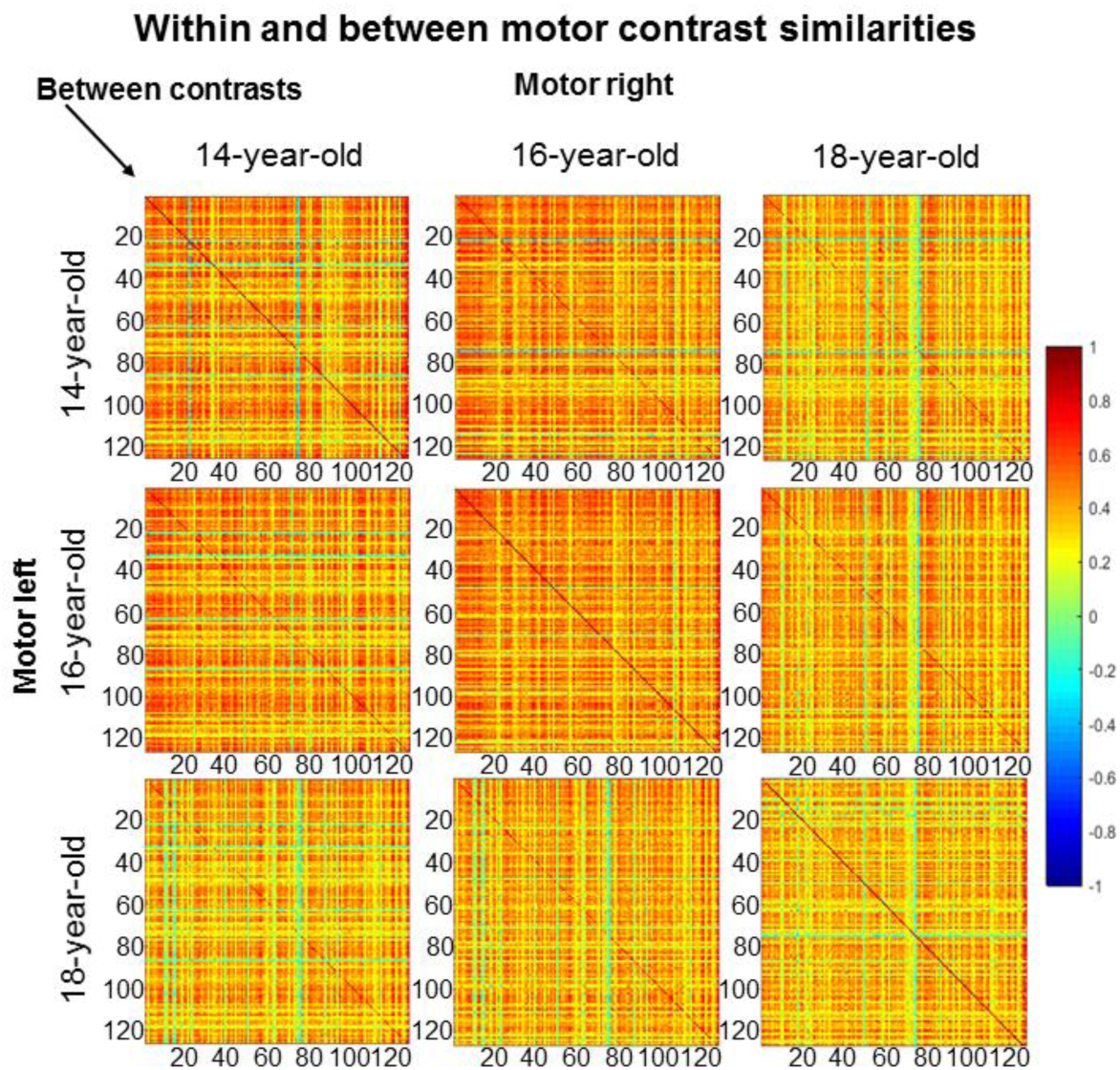
similarity matrices for motor contrasts (one contrast for motor response right hand, one contrast for motor response left hand); between-contrast similarities out of the same session can be interpreted as another form of split-half measure

